# A generative modeling approach for interpreting population-level variability in brain structure

**DOI:** 10.1101/2020.06.04.134635

**Authors:** Ran Liu, Cem Subakan, Aishwarya H. Balwani, Jennifer Whitesell, Julie Harris, Sanmi Koyejo, Eva Dyer

**Affiliations:** School of Electrical & Computer Engineering, Georgia Institute of Technology; Montreal Institute for Learning Algorithms, University of Montreal; Neuroanatomy Division, Allen Institute for Brain Science; Computer Science, University of Illinois at Urbana Champaign; Department of Biomedical Engineering, Georgia Institute of Technology

**Keywords:** variational autoencoder, interpretable deep learning, brain architecture, neuroanatomy

## Abstract

Understanding how neural structure varies across individuals is critical for characterizing the effects of disease, learning, and aging on the brain. However, disentangling the different factors that give rise to individual variability is still an outstanding challenge. In this paper, we introduce a deep generative modeling approach to find different modes of variation across many individuals. To do this, we start by training a variational autoencoder on a collection of auto-fluorescence images from a little over 1,700 mouse brains at 25 micron resolution. To then tap into the learned factors and validate the model’s expressiveness, we developed a novel bi-directional technique to interpret the latent space–by making structured perturbations to both, the high-dimensional inputs of the network, as well as the low-dimensional latent variables in its bottleneck. Our results demonstrate that through coupling generative modeling frameworks with structured perturbations, it is possible to probe the latent space to provide insights into the representations of brain structure formed in deep neural networks.

## 1 Introduction

Understanding how disease, learning, or aging impact the structure of the brain is made difficult by the fact that neural structure varies across individuals [14,5]. Thus, there is a need for better ways to model individual variability that provide accurate detection of structural changes when they occur. Traditional approaches for modeling variability [5,4] require extensive domain knowledge to produce handcrafted features e.g., volumetric covariance descriptors over pre-specified regions of interest (ROIs) [19,13]. However, in high-resolution datasets where micron-scale anatomical features can be resolved, it is unclear i) which features best describe changes of interest across many brains, and ii) how to extract these features directly from images. Thus, unsupervised data-driven solutions for discovering variability across many brains are critical moving forward.

In this work, we introduce a deep learning model and strategy for interpreting population-level variability in high-resolution neuroimaging data (Figure 1). Our model is a regularized variant of the variational autoencoder (VAE) called the *β*-VAE [7,2], and consists of an *encoder* and a *decoder* which work together to first distill complex images into a low dimensional latent space and next, expand this low-dimensional representation to generate high resolution images. Therefore, to gain insight into what the complete model has learned from the data, we take a *bi-directional* approach to characterize how latent components both are impacted by perturbations to specific regions in the input, via the encoder and consequently impact specific regions of the generated output, via the decoder. We provide new strategies for understanding how different brain regions are mapped to latent variables within the network, an important step towards building an interpretable deep learning model that gives insight into how changes in different brain regions may contribute to population-level differences.

**Fig. 1:**
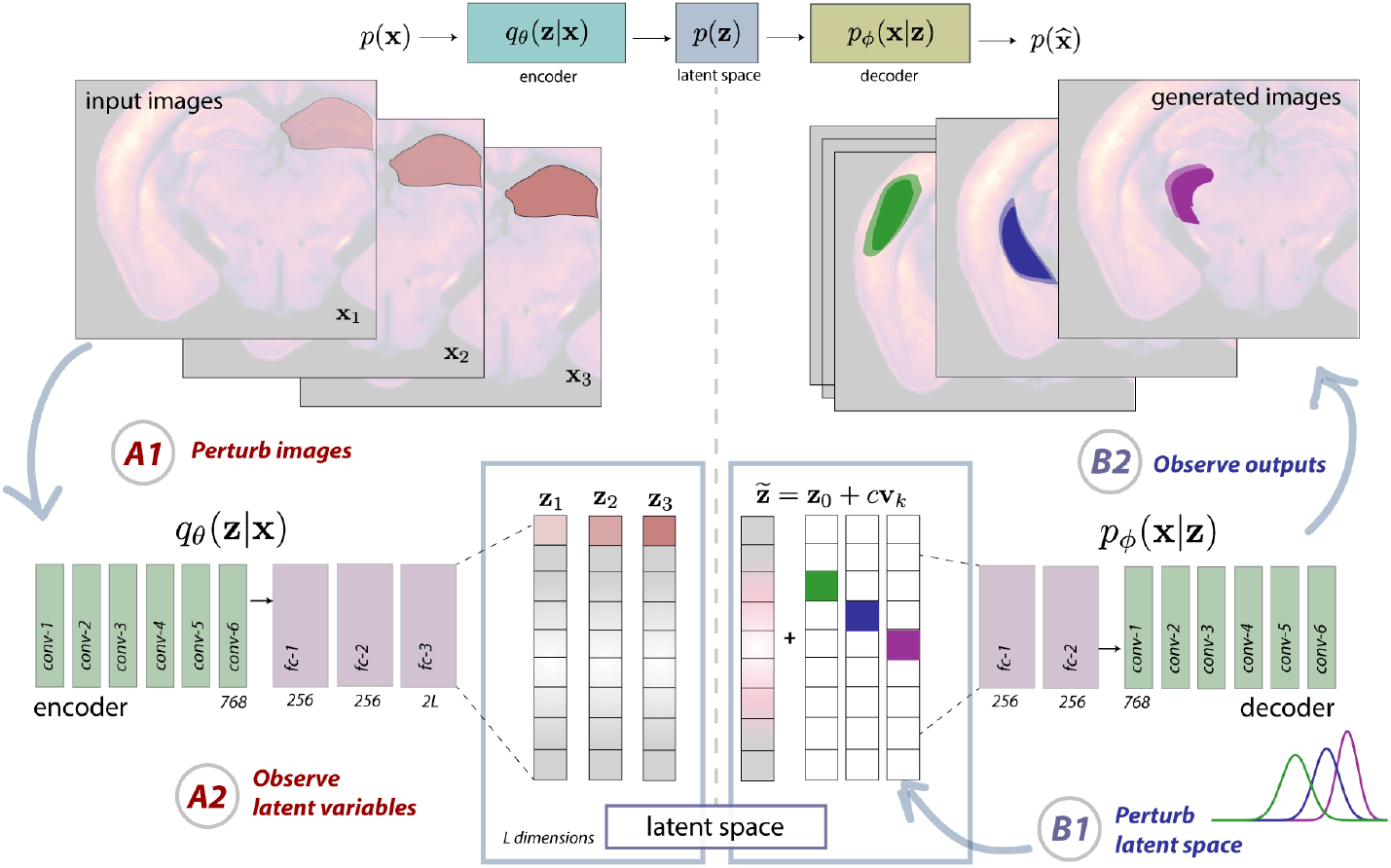
Visualization of our bi-directional approach for analyzing variational autoencoders trained to generate brain imagery. On the left, we show a specific ROI being manipulated in a collection of input images (A1) and how this perturbation might result in a distinct shift in the latent representations (A2) formed from these inputs. On the right, we show the reverse process, where we perturb the latent space (B1) and observe the generated output images (B2). The architecture of the VAE is also depicted, where the latent dimension varies but the rest of the architecture is fixed.

We applied this method to a collection of roughly 1,700 mouse brain images at 25 micron resolution from different individuals in the Allen Mouse Connectivity Atlas. By tuning the regularization strength in the *β*-VAE, we found that it is possible to both generate plausible brain imagery and denoise images corrupted by a number of artifacts. Our investigation into the latent space of this model revealed a number of interesting findings. First, we found that information contained within the latent space is often asymmetric, with artifacts and noise being stored in one direction and biologically meaningful variance observed across many individuals in a separate direction within the same latent factor. Second, we found that multiple units appear to generate outputs that exhibit biologically meaningful variation. Our results demonstrate that the proposed approach can be used to systematically find latent factors that are tuned to specific ROIs, and show that generative modeling approaches can be used to reveal informative components of individual variability.

The contributions of this paper include: (i) the creation and specification of a *β*-VAE that can model high-resolution structural brain images, (ii) a bidirectional approach for revealing relationships between brain regions and latent factors in a deep generative network, and (iii) demonstration that structured perturbations to both image inputs and the latent space can reveal biologically meaningful variability.

## 2 Methods

### 2.1 Model details

Low-dimensional models are used throughout machine learning to represent complex data with only a small set of latent variables. In deep learning, a bottleneck, i.e., layer with small width inside the neural network, often forces a low-dimensional modeling of data. The VAE couples an autoencoder architecture [8,17] with a variational objective, thus providing a probabilistic view towards the generation of new high-dimensional data samples [9,16]. Much like regular autoencoders, VAEs embed information from the image space 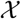 into a latent space 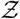 with latent dimension *L* via an encoder, and transform elements from the latent space into those in the image space via a decoder. The relationship between the encoder, decoder, and latent space can be written as:

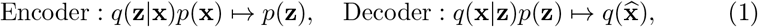

where *p*(**x**) denotes our dataset’s distribution over the high-dimensional image space, *q*(**z**|**x**) and *q*(**x**|**z**) are the distribution of the estimated encoder and estimated decoder respectively, and *p*(**z**) is the assumed prior on latent variables^6^.

To train a good encoder (*θ*) and decoder (*ϕ*), the VAE aims to maximize the following objective:

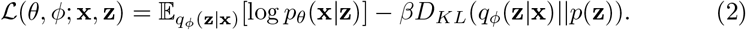

The first term measures the likelihood of the reconstructed samples and the second term measures the KL-divergence between the estimated posterior distribution *q_ϕ_*(**z**|**x**) and the assumed prior distribution. When *β* = 1, the model simplifies to a vanilla-VAE, whereas when *β* is a free parameter, the resulting model is referred to as the *β*-VAE [7]. Increasing the value of *β* encourages a certain degree of clustering, whereas lowering it encourages dispersion of similar elements in the latent space. Thus, by tuning *β* correctly, the model can learn to disentangle latent factors with stronger regularization [7,2].

In our experiments, we used a *β*-VAE with a deep convolutional structure mimicking the DC-GAN architecture [15] (Figure 1). Our encoder had seven convolutional layers followed by three fully connected layers and used the ReLU activation function throughout. The same structure was mirrored for the decoder. The learning rate and batch size were set to 2e-4 and 64 respectively, resulting in a training time of roughly 4 hours on an Nvidia Titan RTX. After performing a grid search (*β* = 1 − 20, *L* = 4 − 20), we selected *L* = 8 and *β* = 3 as our model hyper-parameters since they exhibited performance that was relatively stable (i.e., these parameters produced an inflection point in evaluation metrics). The vanilla VAE’s performance also exhibits an inflection point at the same latent dimension, which further confirmed that this choice holds for different amounts of regularization. In contrast, PCA continues to decrease its approximation error with higher dimensions; however, high-variance artifacts and other sources of noise are very quickly incorporated into the model when the latent dimension increases beyond 30.

### 2.2 Bi-directional latent space analysis

As images in our dataset are spatially aligned to an atlas, understanding how different regions of the pixel space are mapped to latent variables within the network can be a critical first step in building an interpretable model that gives insight into how different brain regions may contribute to population-level differences. To do this, we develop a bi-directional approach to investigate the interaction between the image space and the *β*-VAE’s latent space (see Figure 1). By understanding how the encoder and decoder work together to represent spatial changes in the data, we can build a more informed look into how brain structure can be modeled effectively within deep networks [10,20].

In one direction, we can map a latent variable’s *receptive field* (left, Figure 1), i.e. which pixels in the input space impact each latent factor’s activations. If changing the content of a region of the input image does not impact a specific unit, then the manipulated region is not in the unit’s receptive field. To model this perturbation, let 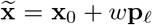 denote the perturbed input image, where **x**_0_ is the original image, **p**_*ℓ*_ is a region specific (spatially localized) perturbation, and *w* is the perturbation weight. By designing these perturbations to examine the responses of the units to changes in specific brain regions of interest, we can study the regional specificity of different units.

In the other direction, we can map a latent variable’s *projective field* (right, Figure 1), or the parts of space that a latent variable affects when a new image is generated. To make this precise, let **v**_*k*_ be a canonical basis vector with a one in the *k*^th^ entry and zeros otherwise, and let *c* denote the interpolation weight. The perturbed latent variable is given by 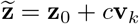. In the simplest case, to visualize these changes, we collapse the generated outputs and measure their per-pixel variance. This provides a measure of which pixels are strongly modified when a specific latent factor changes.

## 3 Results

### 3.1 Dataset and Pre-processing

To build a generative model of brain structure, we utilized a collection of 1,723 registered images from the Allen Institute for Brain Science’s (AIBS) Mouse Connectivity Atlas [12] that is accessible through the Allen Institute’s Python-based SDK [1] (http://connectivity.brain-map.org/). The connectivity atlas consists of 3D image volumes acquired using serial 2-photon tomography (STP) collected from whole mouse brains (0.35 *μ*m x 0.35 *μ*m x 100 *μ*m resolution, 1TB per experiment). Rather than using the fluorescence signal obtained from the viral tracing experiments (green channel), we obtained the auto-fluorescence signal acquired from each of the injected brains (red channel), which captures brain structure and information about overall cell density and axonal projection patterns. Our models were trained on 2D slices extracted from near the middle of the individual brains (slice 286 out of 528) from each of the individuals in our dataset. This particular coronal slice was selected because it reveals key brain areas, including the hippocampus (HPF), regions of thalamus (TH), and parts of striatum (STR) (Figure 2A). The images were then downsampled from 0.35 microns to 25 microns, and centre-cropped to produce an image of size 320 × 448 (input dimension). In order to mitigate the effects of leakage of fluoresence signal, we pre-processed the data by adjusting image brightness to the dataset’s average brightness and removing the resulted outliers.

**Fig. 2:**
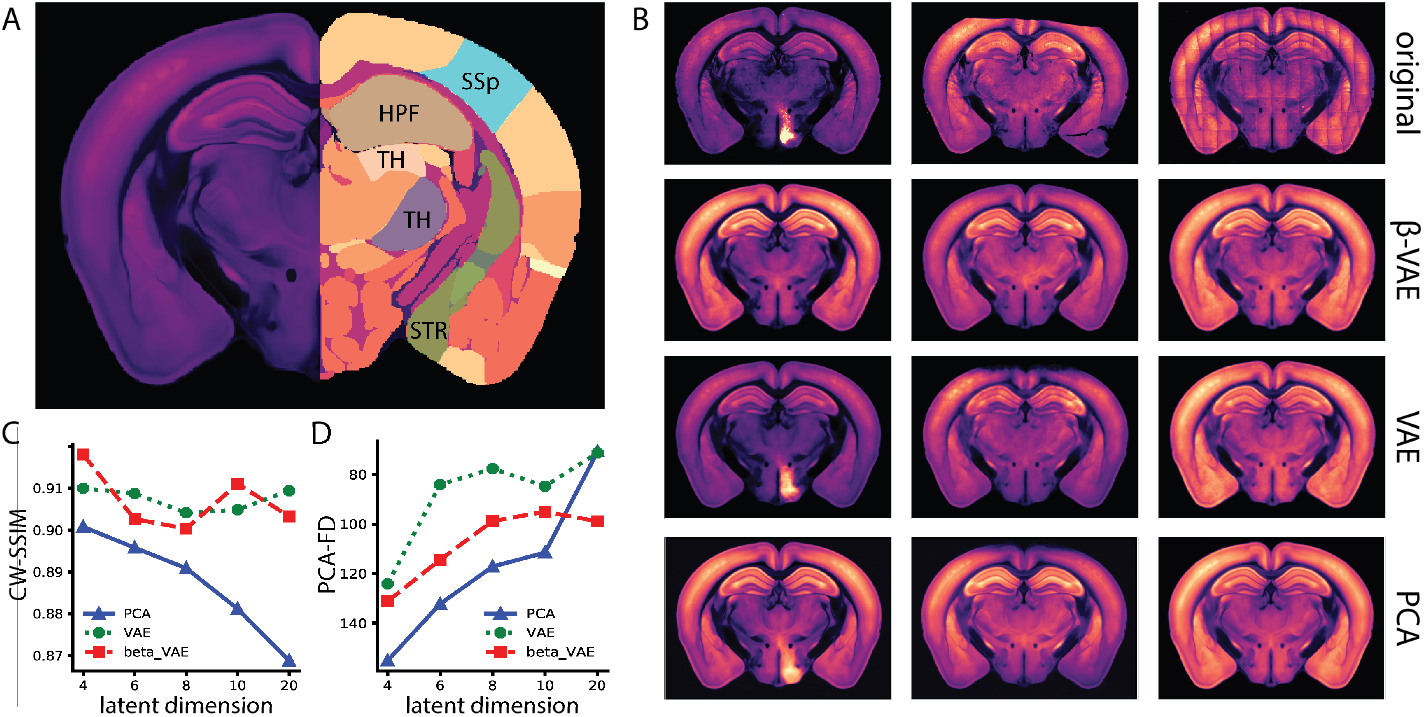
Evaluation of image synthesis and denoising performance. **(A)** The left side shows the average template of brain slice and the right side shows the atlas of the interested regions of brain structures, including somatosensory areas (SSp), hippocampal formation (HPF), striatum (STR), and parts of the thalamus (TH). **(B)** We show examples of images with different types of artifacts and the reconstructions obtained with all three models. The first column shows the corruption artifacts, the second column shows the physical breakage artifacts, and the third column shows the grid-like artifacts. These types of artifacts are representative in real-world dataset. The CW-SSIM and PCA-based FD scores for all three models are compared in **(C)** and **(D)**, respectively.

### 3.2 Evaluations and comparisons

To evaluate the image generation capability of our *β*-VAE, we compared its performance qualitatively and quantitatively with a vanilla-VAE and PCA. We first sought to examine the reconstruction and denoising properties of the models by seeing how they performed when supplied with images containing three different types of artifacts: (i) corrupted bright areas due to leakage from the green channel, (ii) physical sectioning artifacts (missing data), and (iii) grid-like artifacts from scanning (Figure 2B, Supp. Materials S1). In these and other examples, we found that the *β*-VAE did the best job of removing artifacts from data. This ability of the *β*-VAE is particularly pronounced in the case of class (i, ii) artifacts, where both PCA and VAE fail to reject the signal leaking into the channel of interest and fail to recover missing data respectively. We observe that the *β*-VAE tends to learn a more accurate distribution over the dataset, while the vanilla-VAE overfits to the noise, and PCA given its linear nature, does not deviate much from the mean in terms of its structural details.

To quantify the quality of reconstructed images and investigate how changing the latent dimension effects the the model performance, we computed the complex wavelet structural similarity (CW-SSIM) [18] and the PCA-based Frechet distance (PCA-FD) [6,11] for all three models as we varied the number of latent variables. We found that after tuning the model’s latent dimension, the *β*-VAE (*L* = 8) outperforms both the vanilla-VAE and PCA on our chosen metrics (Figure 2 C,D). Analysis of the CW-SSIM scores suggests that PCA was unable to capture high-dimensional textural details and high-frequency components of brain images for low dimensions and then as the dimension increases it quickly starts to represent artifacts in data. The PCA-FD scores on the other hand suggest that the VAE models capture more variability across the data and better matches the overal global distribution. But it does so at the expense of overfitting to the dataset and generates more noisy realizations. However, the *β*-VAE appears to successfully capture variability without reconstructing artifacts. It is also worth noting that when the latent dimension is very large, the performance of the VAE drops significantly, in line with previous observations of how VAEs fail in higher dimensions [20]. These results provide initial evidence that regularization is helpful for denoising the wide variety of artifacts, while also capturing the data’s distributional properties.

### 3.3 Interpreting the latent factors

After confirming that our model can generate high quality images and denoise data, we next explored its interpretability with the bi-directional analysis method described in Section 2.2 (Figure 1).

We first examined the latents’ projective field, by generating a collection of images via a dense uniform interpolation of the latent space, varying a single latent factor at a time (*c* = ±7, with a step size of 0.005). When we examined the resulting images, in many of the factors, we observed that localized bright noise artifacts (type i) were essentially synthesized at the extrema of the interpolation space. Interestingly, we observed asymmetries in this representational strategy: Type (i) artifacts, while not usually recovered by the decoder, were more likely recapitulated when moving far into the space of negative interpolation weights (Supp. Material S4). This indicates that a data example is encoded in the left tail of the distribution, centered about the original sample. On the other extreme, in high positive interpolation weights, we observed some high variance regions, but they all aligned with patterns of variability that we know are useful for encoding biologically meaningful variance in the whole dataset. We thus divided the interpolation space into three groups: (i) small negative weights, (ii) small positive weights, and (iii) large weights. All three groups of images were compressed by computing their pixel-level variances and stacking them into different channels of an RGB color image; thus allowing for the visualization of the impact of all three types of perturbations to the latent space, on the image domain jointly (Figure 3A). When we examined the variability heatmaps, we observed that factors 2, 6, and 8 (marked with stars in Figure 3A) recapitulated structures present due to outliers. As mentioned before, these noise artifacts appear to be stored in parts of the latent space that are accessed when we apply large negative interpolation weights to true examples (visualized in the stratified variance map). In the other parts of this interpolation space, we found biologically meaningful variance, where the model highlights key ROIs including the barrel fields of somatosensory cortex, hippocampus, and retrosplenial areas in cortex. These results provide initial evidence that VAE models can be used to decompose variability in complex data with different types of noise and artifacts.

**Fig. 3:**
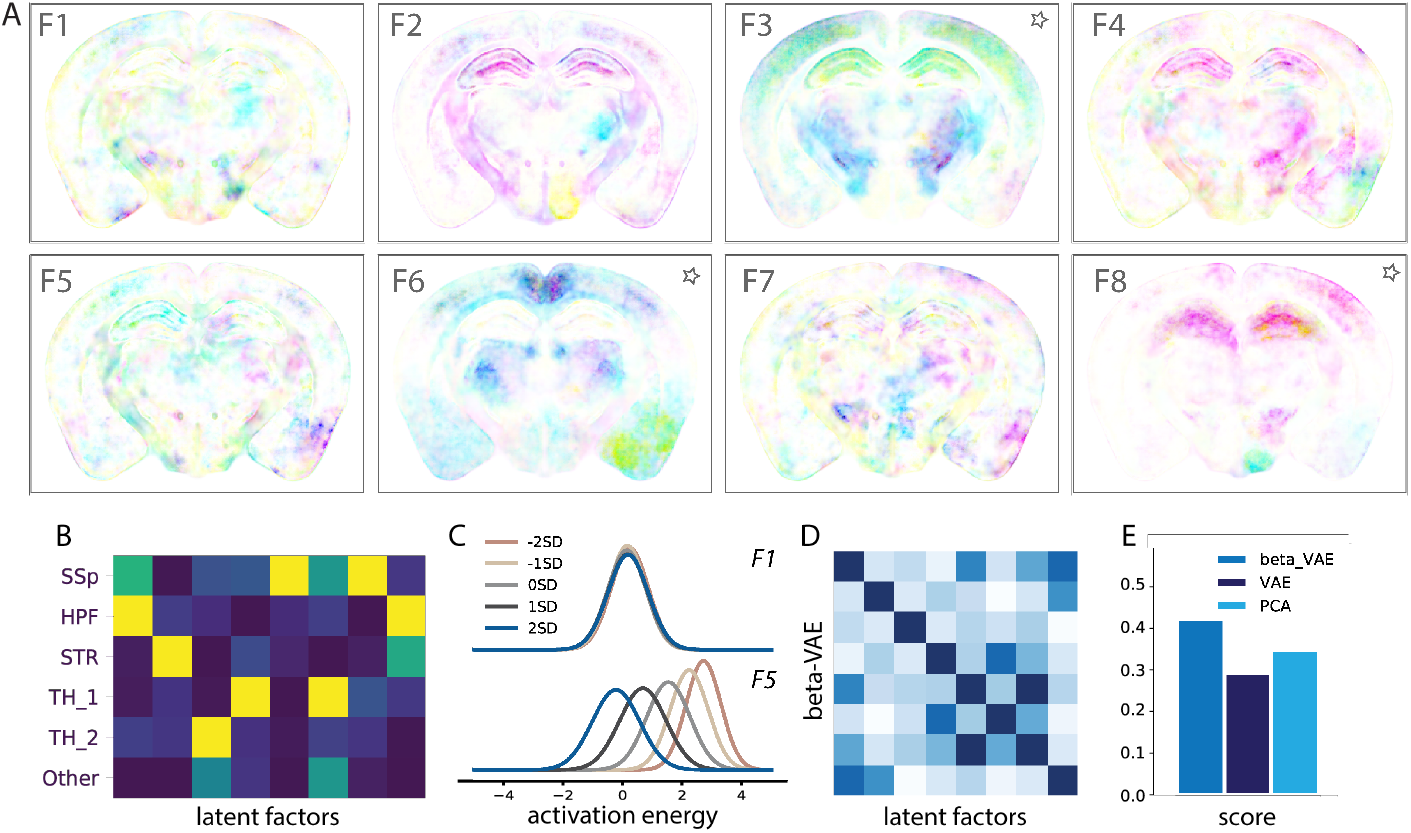
Model interpretability. **(A)** Stratified pixel-level variance heatmaps, where cyan and magenta represent variance generated from interpolation weights in the quartile above average and below average, respectively, and yellow represents points with high interpolation weights. **(B)** Each entry of this matrix contains the KL-divergence between a specific latent factor’s activation distribution when perturbing a specific brain ROI. For each ROI, the KL-divergence across all eight latent dimensions is normalized and their relative impact are displayed in color (blue is low impact, yellow is high impact). **(C)** We show how perturbing the image brightness in HPF region impacts the activation distribution for two factors (F1, F5). **(D)** The covariance between the impact across factors provides a visualization of the similarity between how different latent variables impact specific ROIs. **(E)** We score the *β*-VAE, VAE, and PCA models by measuring how uncorrelated two latent factors are in terms of how they are impacted by ROI-based perturbations. The *β*-VAE model has the highest score, suggesting that it has the most uncorrelated representation of the tested ROI-specific perturbations.

After exploring the projective field of the units in the latent space, we next asked whether we could understand properties about each unit’s *receptive field.* To do so, we selected a set of high-quality images without obvious artifacts, applied masks to remove all content from different ROIs, modulated their intensity with perturbation weights *w*, and fed these perturbed images into the encoder (Supp. Material S2). We then fit a Gaussian to the resulting latent codes across all image examples (*n* = 832) (Figure 3C, Supp. Materials S3). The results of this perturbation analysis revealed multiple units that are not modulated by certain ROIs, and also that many factors have spatially localized receptive fields. We found that perturbations to the hippocampus (HPF) impacted almost all of the latent variables, and striatum also has wide reaching impacts. This seems to align with the fact that variability in these areas is more complex and thus it is necessary to encode this variance over multiple factors. We computed the KL-divergence between the activation distributions for two extreme brightness values, and then compiled all these distances into a 6×8 matrix (Figure 3B). This matrix quantifies the impact that missing information from a ROI has on activations in each latent variable in the model. One interesting result from our analysis is that, in some cases, the receptive field and projective field may not be spatially aligned (see Factor 8, HPF). Our results reveal that receptive and projective fields can be asymmetric, and thus it is critical to map input-output relationships from the image space to the latent space and back again.

## 4 Discussion

In this paper, we have developed a methodology to interpret the population-level variability in structural brain images in deep generative models. We introduced a method for visualizing the impact of perturbations to the latent space of a *β*-VAE and used it to dissect models trained on brain imagery from many individuals.

In our current study, we used a *β*-VAE model because of its simplicity and flexibility. However, there are other interpretable VAE variants that have been proposed to facilitate disentanglement [2,20,3]; these methods incorporate additional priors to encourage separability across latent dimensions. As our approach is relatively general, exploring deep representations in the way described could provide a way to visualize the representations formed in generative models for other applications in medical imaging. With new strategies for interpreting how deep networks represent data, we may be able to develop new regularization strategies to disentangle and interpret population-level variability.

## Acknowledgements

This work was supported by NSF award number IIS-1755871 (ELD, AHB); ELD was also supported by an Alfred P. Sloan Fellowship and NIH Award No. 1R24MH114799-01.

## Supplementary Materials

**Fig. S1:**
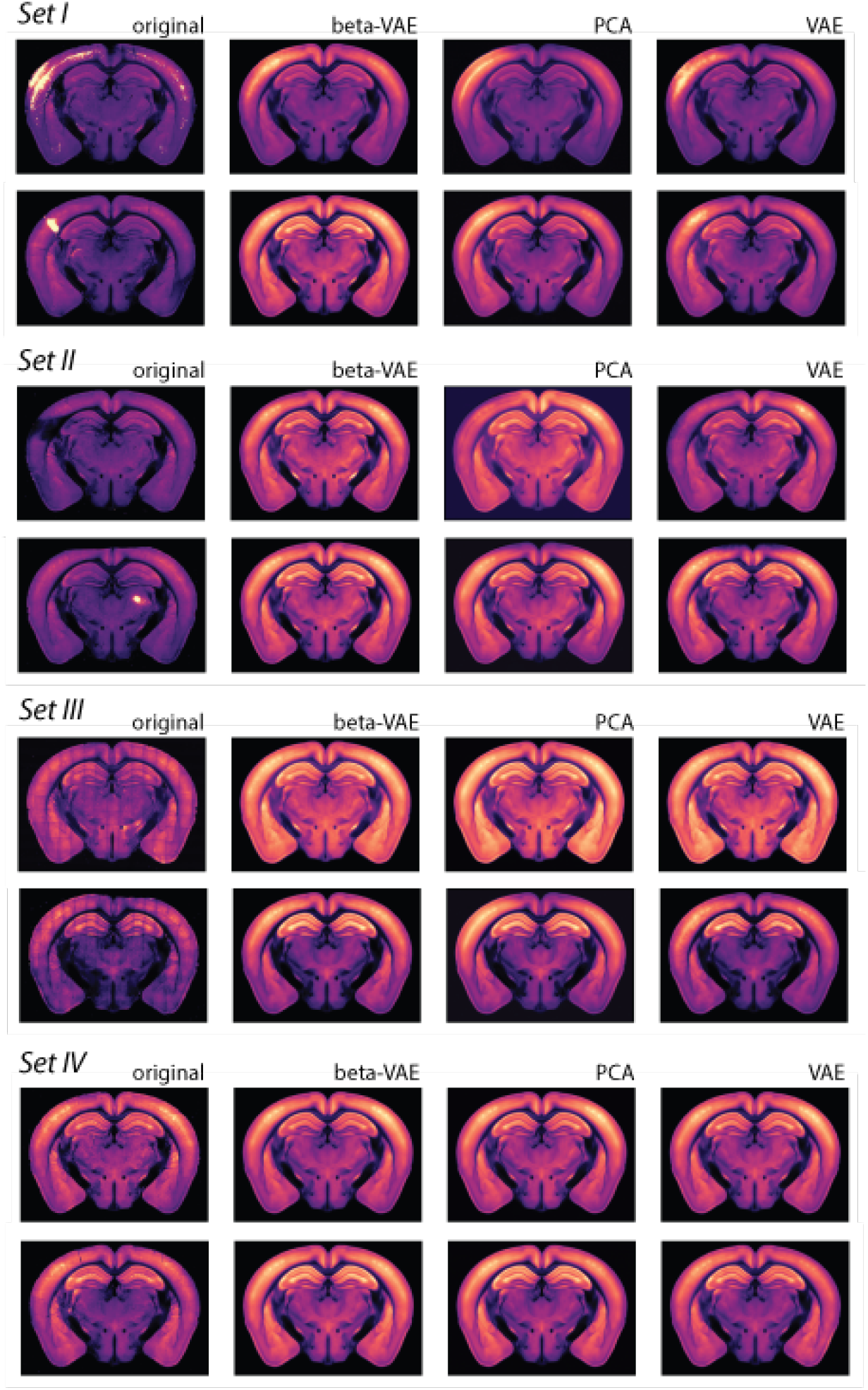
Additional examples of model comparison. Similar as Figure 2B where different types of artifacts are fed into three models.

**Fig. S2:**
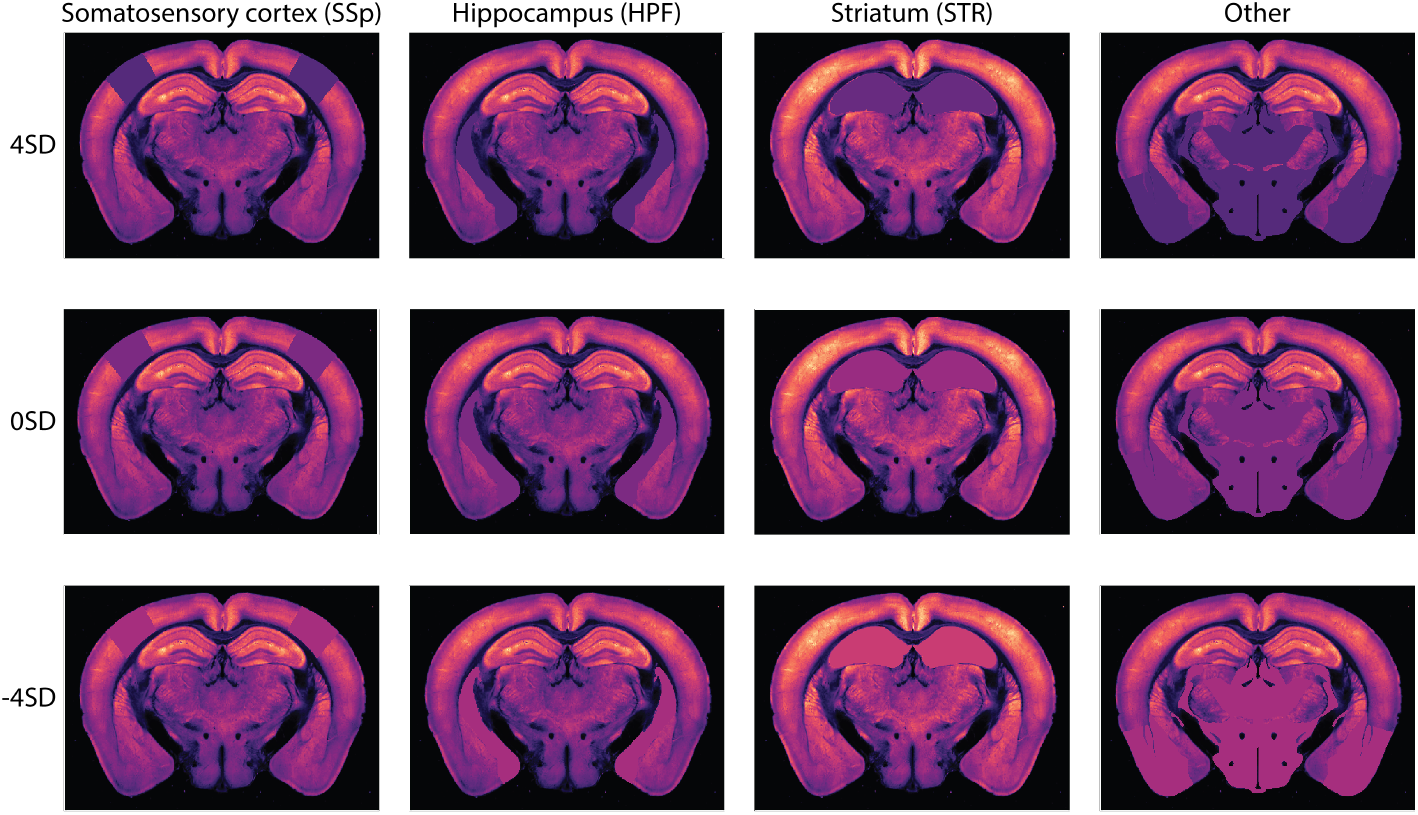
Examples of masked images used to probe the latent space. Different brightness levels of different ROIs are calculated specifically to mask the original images.

**Fig. S3:**
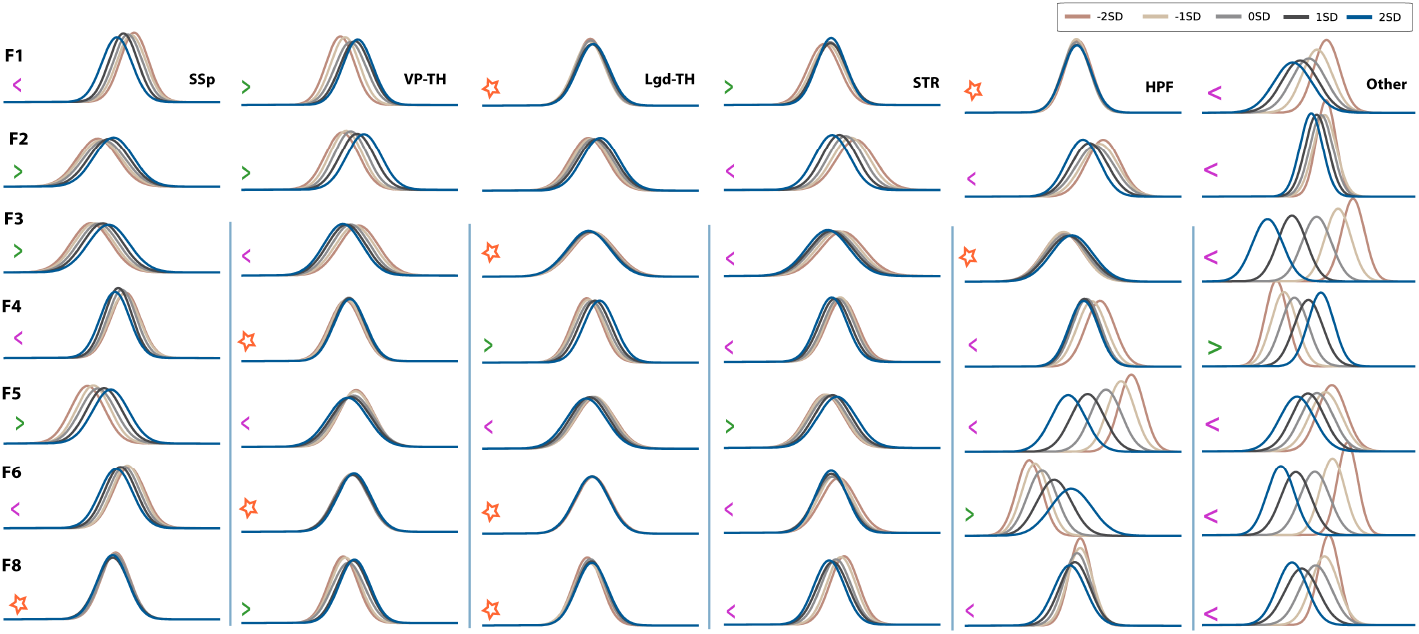
Results of image perturbation experiment. Each row represents a different latent factor and each column represents a specific ROI. A complete version of Figure 3C.

**Fig. S4:**
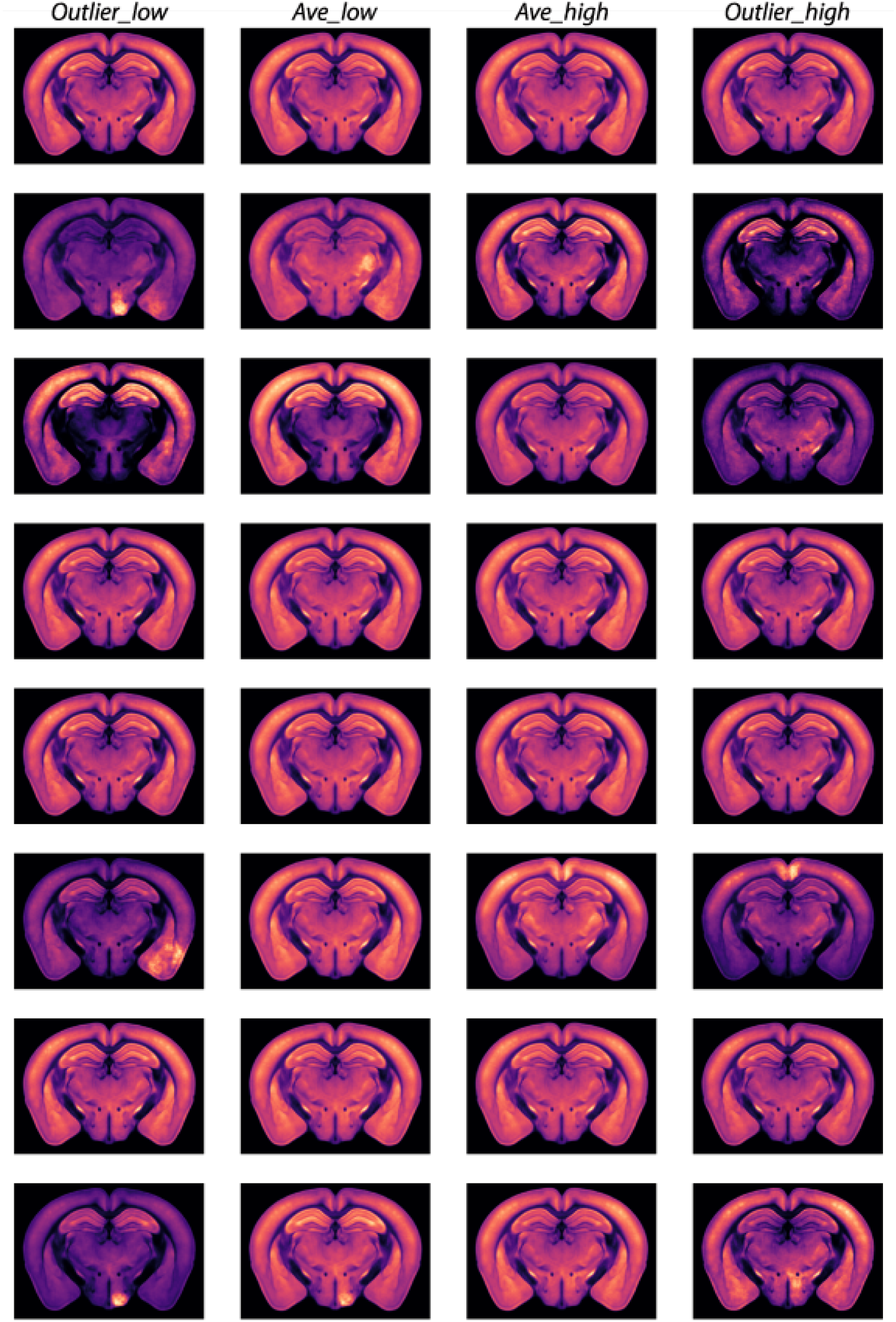
Examples of output images generated with latent interpolation. They are used to produce the heatmaps in Figure 3A.

## Footnotes

6 For simplicity, the prior is typically assumed to be Gaussian, 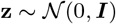.

